# Single cell RNA sequencing reveals differential cell cycle activity in key cell populations during nephrogenesis

**DOI:** 10.1101/2020.09.16.300293

**Authors:** Abha S. Bais, Débora M. Cerqueira, Andrew Clugston, Jacqueline Ho, Dennis Kostka

## Abstract

The kidney is a complex organ composed of more than 30 terminally differentiated cell types that all are required to perform its numerous homeostatic functions. Defects in kidney development are a significant cause of chronic kidney disease in children, which can lead to kidney failure that can only be treated by transplant or dialysis. A better understanding of molecular mechanisms that drive kidney development is important for designing strategies to enhance renal repair and regeneration. In this study, we profiled gene expression in the developing mouse kidney at embryonic day 14.5 at single cell resolution. Consistent with previous studies, clusters with distinct transcriptional signatures clearly identify major compartments and cell types of the developing kidney. Cell cycle activity distinguishes between the “primed” and “self-renewing” sub-populations of nephron progenitors, with increased expression of the cell cycle related genes *Birc5, Cdca3, Smc2* and *Smc4* in “primed” nephron progenitors. Augmented *Birc5* expression was also detected in immature distal tubules and a sub-set of ureteric bud cells, suggesting that *Birc5* might be a novel key molecule required for early events of nephron patterning and tubular fusion between the distal nephron and the collecting duct epithelia.

## INTRODUCTION

The mammalian kidney has evolved to provide critical adaptive regulatory mechanisms, such as the excretion of waste, and the maintenance of water, electrolyte and acid-base homeostasis to the body. These functions require the coordinate development of specific cell types within a precise three-dimensional pattern. Defects in kidney development are amongst the most common malformations at birth. Congenital anomalies of the kidney and urinary tract (CAKUTs) represent more than 20 percent of birth defects overall [1], and they account for a large fraction of chronic kidney disease and renal failure in children [2]. For example, the number of nephrons formed at birth is thought to be an important determinant of renal function, because reduced nephron numbers are often observed in humans with primary hypertension and chronic kidney disease [3, 4]. An estimated 37 million people in the United States (∼15% of the population) have chronic kidney disease (CKD) [5, 6] that can lead to kidney failure requiring transplant or dialysis. Development of strategies to enhance renal repair or regeneration are needed to reduce the morbidity and mortality associated with kidney disease, and they are dependent on a better understanding of the molecular genetic processes that govern kidney development.

Nephrons form the functional units of the kidney and are derived from a nephron progenitor (NP) cell population, also known as cap mesenchyme. These cells are capable of self-renewal, which is necessary to generate an appropriate number of nephrons during the course of embryogenesis and development. They are also multipotent, that is they have the ability to differentiate into the multiple cell types of the mature nephron [7, 8]. More specifically, multipotent Cbp/P300-Interacting Transactivator 1 (Cited1)-positive/ Sine Oculis Homeobox Homolog 2 (Six2)-positive nephron progenitors give rise to multiple nephron segments, and are termed “self-renewing” nephron progenitors [9]. The transition of nephron progenitors into epithelialized structures is dictated by a series of tightly orchestrated signaling events. Of this, Bone morphogenetic protein 7 (Bmp7) induces the initial exit of Cited1^+^/Six2^+^ cells into a Cited1^-^/Six2^+^ state, which marks nephron progenitors “primed” for differentiation by ureteric bud-derived Wnt family member 9b (Wnt9b)/β-catenin signaling. Conversely, remaining Cited1^+^/Six2^+^ nephron progenitors are kept in an undifferentiated and self-renewing state in response to Fibroblast growth factor 9 (FGF9), Wnt and BMP7 signals [10-18].

Upon Wnt9b/β-catenin stimulation, nephron progenitors undergo a mesenchymal to epithelial transition to form pre-tubular aggregates, which then proceed to develop sequentially into polarized epithelial renal vesicles, comma- and then S-shaped body structures. Cells in the proximal portion of the S-shaped body differentiate into podocytes (glomerular development), while its mid- and distal portions give rise to tubular segments of the nephron, which are subdivided into proximal tubules, loops of Henle and distal tubules [19], **Figure 1A**. During the S-shaped stage of glomerular development, developing podocytes secrete vascular endothelial growth factor A (VEGF-A), which attracts invading endothelial cells into the cleft of the S-shaped body. Platelet-derived growth factor-β (PDGFβ) signal produced by endothelial cells mediates the recruitment of mesangial cells, which invade the developing glomerulus and attach to the forming blood vessels. By the end of maturation, the glomerulus consists of four specified cell types: the fenestrated endothelium, mesangial cells, podocytes and parietal epithelial cells of the Bowman’s capsule [20-24].

**Figure 1:**
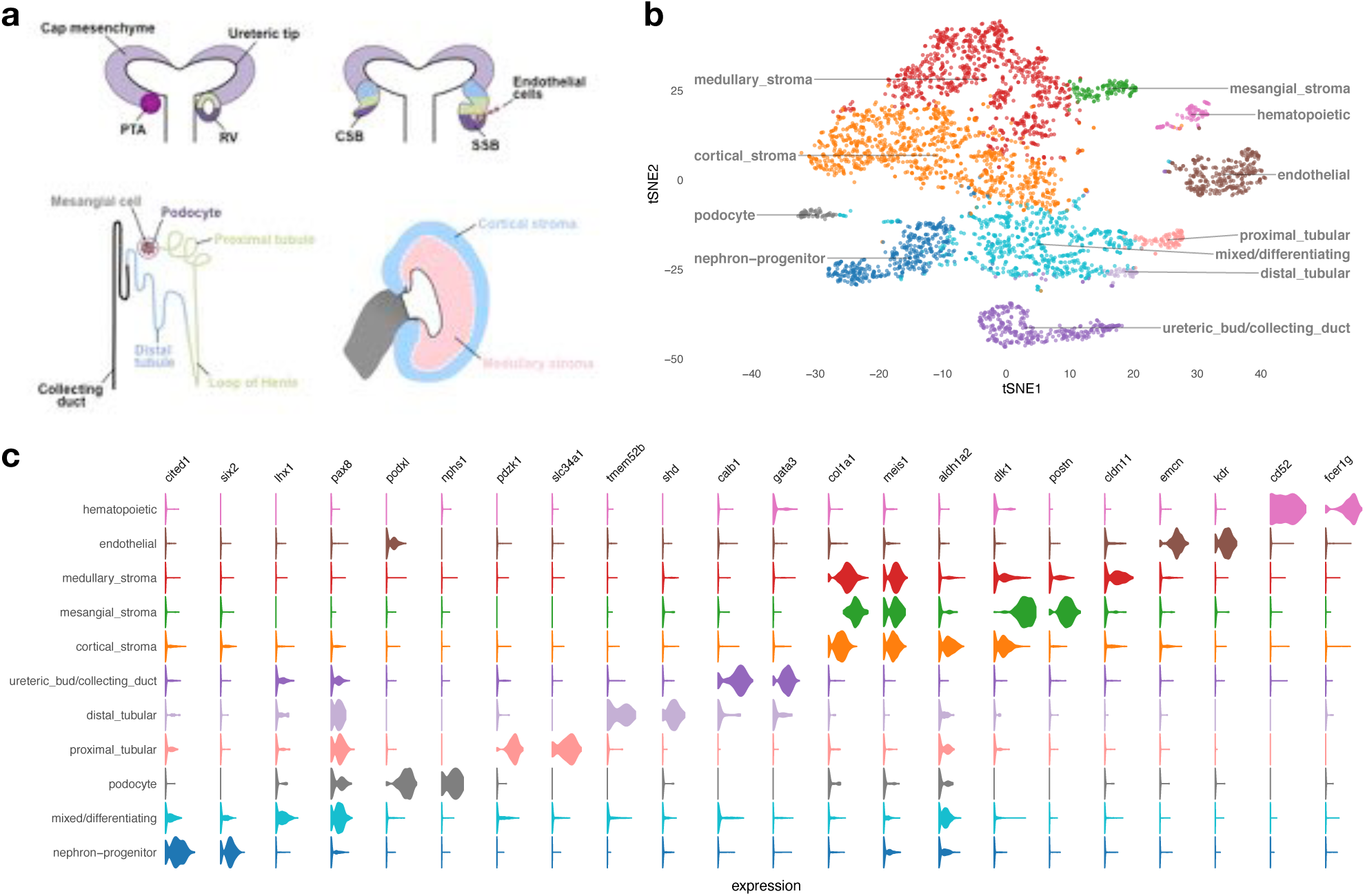
Developing embryonic day 14.5 mouse kidney cell types. ***(a)*** Schematic illustration of nephron induction and patterning. In response to signals from the ureteric bud, the metanephric mesenchyme condenses and forms a cap of nephron progenitors (=cap mesenchyme) around the ureteric bud tips. Next, a sub-population of nephron progenitors undergoes a mesenchymal to epithelial transition to form pre-tubular aggregates (PTA), which develop sequentially into renal vesicles (RV), comma-shaped body (CSB) and S-shaped body (SSB). Endothelial cells are attracted into the cleft of the SSB. Color-coded map indicates the cell fate relationship of progenitor regions in SSB structure (upper right) and adult nephron structure (lower left). Schematic of a lateral view of the metanephric kidney depicting the cortical and medullary stroma (lower right). ***(b)*** tSNE plot showing the eleven cell clusters in the embryonic mouse kidney, with cell clusters corresponding to major components indicated by color. ***(c)*** Violin plots of gene expression for known lineage-associated genes (columns), stratified by cluster (rows). Our data clearly identifies cells from the major structural components of the developing kidney.

Single cell RNA sequencing (scRNA-seq) technology offers the ability to comprehensively identify the transcriptional and (inferred) cellular composition of the developing kidney. Recent studies in developing mouse [25-30] and human kidneys [31-33] have contributed to our understanding of subpopulations of nephron progenitors and stromal cells, lineage fidelity, novel receptor-ligand pathways, and differences between mouse and human kidney development. scRNA-seq also has the potential to inform improvements in our ability to culture nephron progenitor cells [34-36] and produce higher-fidelity human kidney organoids [37, 38], and to develop novel strategies for enhancing renal repair and regeneration.

In this study, we used scRNA-seq to interrogate cell types and transcriptomes within 4,183 cells from one kidney pair of an E14.5 female mouse embryo. Clustering identified eleven clusters corresponding to the major components/cell-types of the developing kidney and revealed expression of known lineage markers in unexpected cell types (e.g., renal stromal markers in nephron progenitors). Pseudotime analysis was utilized to describe transcriptional dynamics as nephron progenitors differentiate. Notably, we find that cell cycle activity distinguishes between the “primed” and “self-renewing” sub-populations of nephron progenitors, with increased levels of the cell cycle related genes *Birc5, Cdca3, Smc2* and *Smc4* in the “primed” sub-population. Moreover, increased *Birc5* expression was also observed in immature distal tubules and in a sub-set of ureteric bud cells, suggesting its involvement in the process of fusion between the distal nephron and the collecting duct epithelia.

## RESULTS

### Single cell gene expression identifies anatomical structures and cell lineages in the developing kidney

New nephrons are induced in response to signals from the ureteric bud throughout nephrogenesis until approximately postnatal day 3 in mice [39]. Thus, we chose to perform scRNA-seq at embryonic day 14.5 (E14.5), a time point at which there is active nephron induction and varying degrees of nephron maturation, to comprehensively interrogate single cell transcriptomes spanning different stages of differentiation during kidney development at mid-gestation. Using one kidney pair from an E14.5 female mouse embryo processed using the 10X Chromium platform and Illumina sequencing, our dataset consists of 4,183 high-quality kidney cells, with a median number of 2,789 genes detected per cell. Grouping cells into eleven clusters (see Methods) reveals major components/cell-types of the developing kidney (**Figure 1a-c, Supplemental Figure 1 and Supplemental Table 1)**. Clusters and key markers are consistent with prior single cell analyses of the developing mouse kidney [25-29]. We observe clear separation of cells of the hematopoietic (*Cd52, Fcer1g*), ureteric bud/collecting duct (*Calb1, Gata3*), and endothelial (*Emcn, Kdr*) lineages from other cells of the developing kidney (nephron progenitors, mixed/differentiating cells, podocytes, tubular cells and stromal cells). Stromal lineages are marked by expression of *Col1a1* and *Meis1*, while cells derived from the nephron progenitor lineage express established marker genes associated with progressive stages of nephron differentiation. Thus, *Cited1* and *Six2* identify nephron progenitors, *Lhx1* and *Pax8* mark mixed/differentiating cells, *Fxyd2* and *Hnf4a* mark tubular cells, and podocytes are marked by *Podxl* and *Nphs1*.

**Table 1:** Co-expression of stromal marker genes and nephron progenitor marker genes in cortical/stromal cells. The cs_gene and np_gene columns show the cortical/stromal and nephron-progenitor marker genes, respectively; the #cells column shows the number of cells expressing both genes in the stromal/cortical cluster (which contains 1,085 cells overall), while the odds_ratio and p_value columns contain odds ratio and p_value of a corresponding Fisher exact test.

| cs_gene | np_gene | #cells | odds_ratio | p_value |
| --- | --- | --- | --- | --- |
| col1a1 | cited1 | 237 | 8.32 | 8.29E-13 |
| col1a1 | crym | 260 | 5.87 | 1.27E-11 |
| col1a1 | gdnf | 407 | 1.73 | 1.87E-03 |
| col1a1 | six2 | 237 | 4.71 | 3.00E-09 |
| meis1 | cited1 | 214 | 2.14 | 2.12E-04 |
| meis1 | crym | 238 | 2.21 | 4.33E-05 |
| meis1 | gdnf | 402 | 2.39 | 5.94E-08 |
| meis1 | six2 | 221 | 2.42 | 1.64E-05 |

Consistent with other reports, we identify three different stromal clusters: medullary stroma (*Col1a1, Meis1* and *Cldn11*), cortical stroma (*Col1a1, Meis1, Aldh1a2* and *Dlk1)* and mesangial stroma (*Col1a1, Meis1, Dlk1, Postn*) [25, 28]. Analyses of *in situ* hybridization data at E14.5 from other reports, as well as GUDMAP (The GenitoUrinary Development Molecular Anatomy Project) and Eurexpress public resources facilitated identification and assignment of these clusters [27, 40-42].

Taken together, these results show that our scRNA-seq data successfully captured major cell types that are expected to be present in the developing kidney at E14.5, including progenitor cells and their derivatives as well as mature cell populations.

### Stratification of cell-types in the nephron progenitor lineage

Next we focused on nephron progenitor cells and their descendant/derived cell types (mixed/differentiating, podocytes and tubular cells). Selecting those cell-types yielded 1,727 cells for further analysis. We are able to clearly distinguish between proximal and distal tubular cells and podocytes, and pseudotime analysis allows us to assess the level of lineage commitment (**Figure 2a**). Nephron progenitor cells (marked by *Six2* and *Cited1* expression) clearly separate into two sub-groups, “self-renewing” and “primed” (**Figure 2b**), see below for more details. Mixed/differentiating cells express transcription factors like *Pax8* and *Lhx1*, which are associated with nephron development and encompass nephron progenitor cells differentiating into tubular cells and podocytes. We note heterogeneity in the mixed/differentiating cell cluster, which likely contains cells with different degrees of differentiation, like pre-tubular aggregate, renal vesicle, comma-, and S-shaped bodies. Pseudotime analysis on this sub-set of cells reconstructs three lineages: differentiation into podocytes, and into proximal and distal tubular cells (**Supplemental Figure 2**). This enabled us to distinguish between mature and immature podocytes and proximal/distal tubular cells (**Figure 2**, see **Methods**). Overall, our data clearly shows the major differentiation trajectories of nephron progenitor cells. For these three lineages, podocytes are marked by increasing expression of *Podxl* and *Nphs1* [43], proximal tubular cells by *Pdzk1* and *Slc34a1* [44], and distal tubular cells by *Tmem52b* and *Shd* [29] (**Supplemental Figure 3** shows a heatmap of genes with pronounced expression differences during nephron progenitor cell differentiation and **Supplemental Table 2** summarized differentially expressed genes).

**Table 2:**
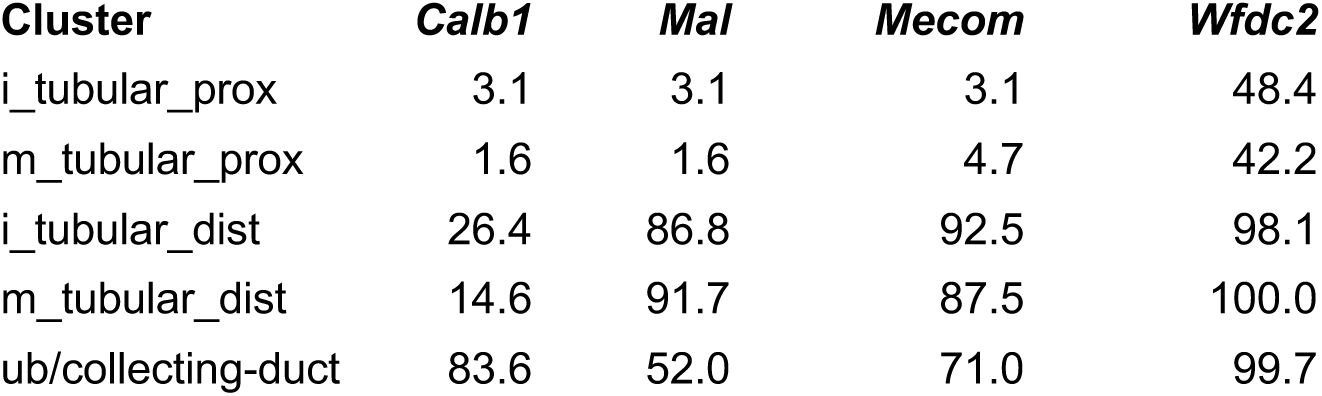
Ureteric bud / collecting duct lineage genes are expressed in distal tubular cells. Shown is the percentage of cells expressing ureteric bud / collecting duct (ub/cd) lineage marker genes *Calb1, Mal, Mecom* and *Wfdc2* for cells from immature and mature distal and proximal tubular clusters, and also from the ub/cd cluster. We see that in the tubular distal lineage, in contrast to the proximal lineage, a significant fraction of cells expresses these ub/cd marker genes.

**Figure 2:**
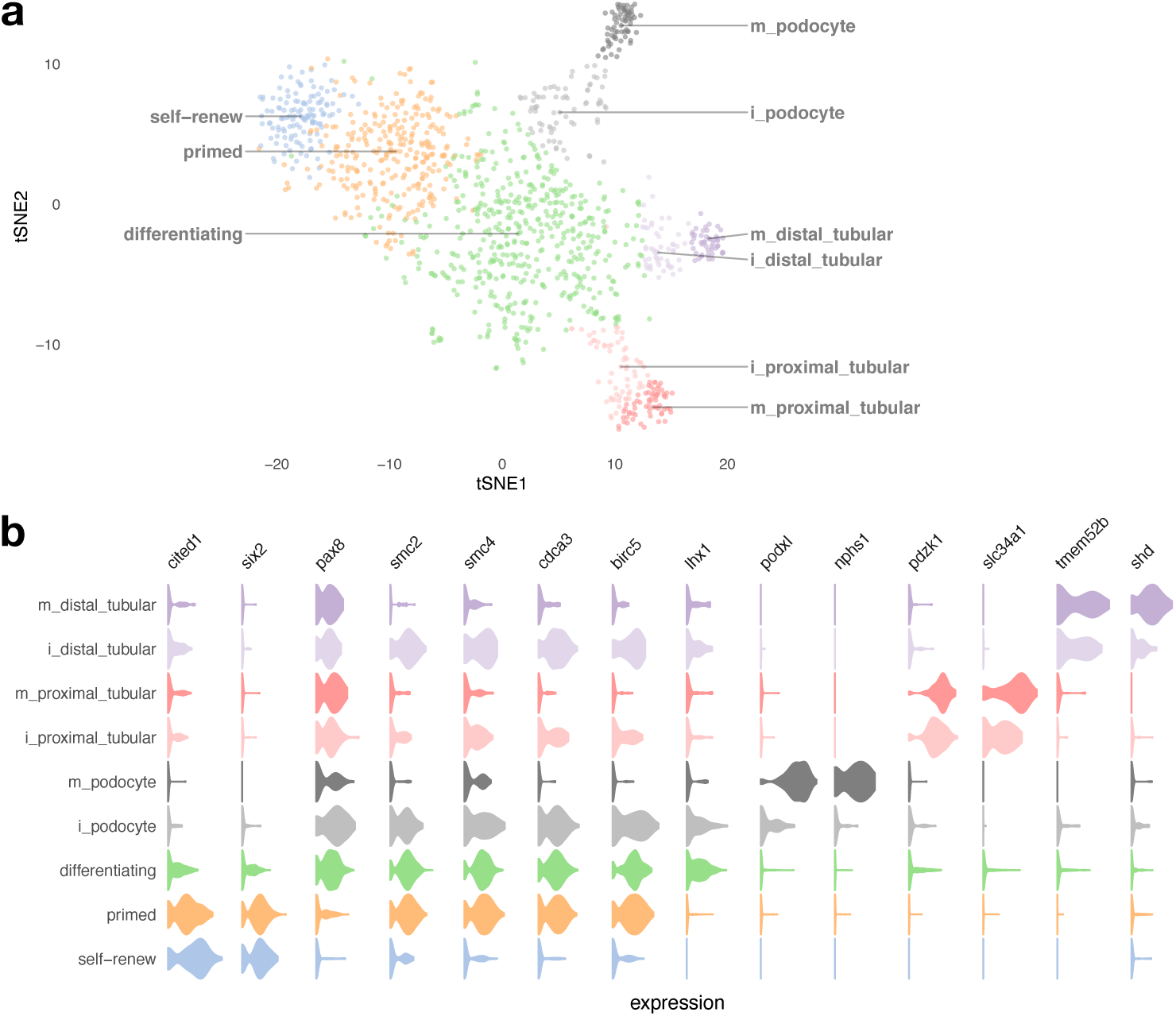
Cell types of the nephron progenitor lineage. Panel ***(a)*** shows a tSNE plot of NP-derived cells, with clusters corresponding to cell types annotated in colors. The prefix “i_” indicates immature cells, while “m_” indicates mature cells. Panel ***(b)*** shows violin plots of gene expression for known lineage-associated genes (columns), stratified by cluster (rows). We observe two types of NP cells (“self-renewing” and “primed”) and clear separation of distal and proximal tubular cells and podocytes in our data.

**Figure 3:**
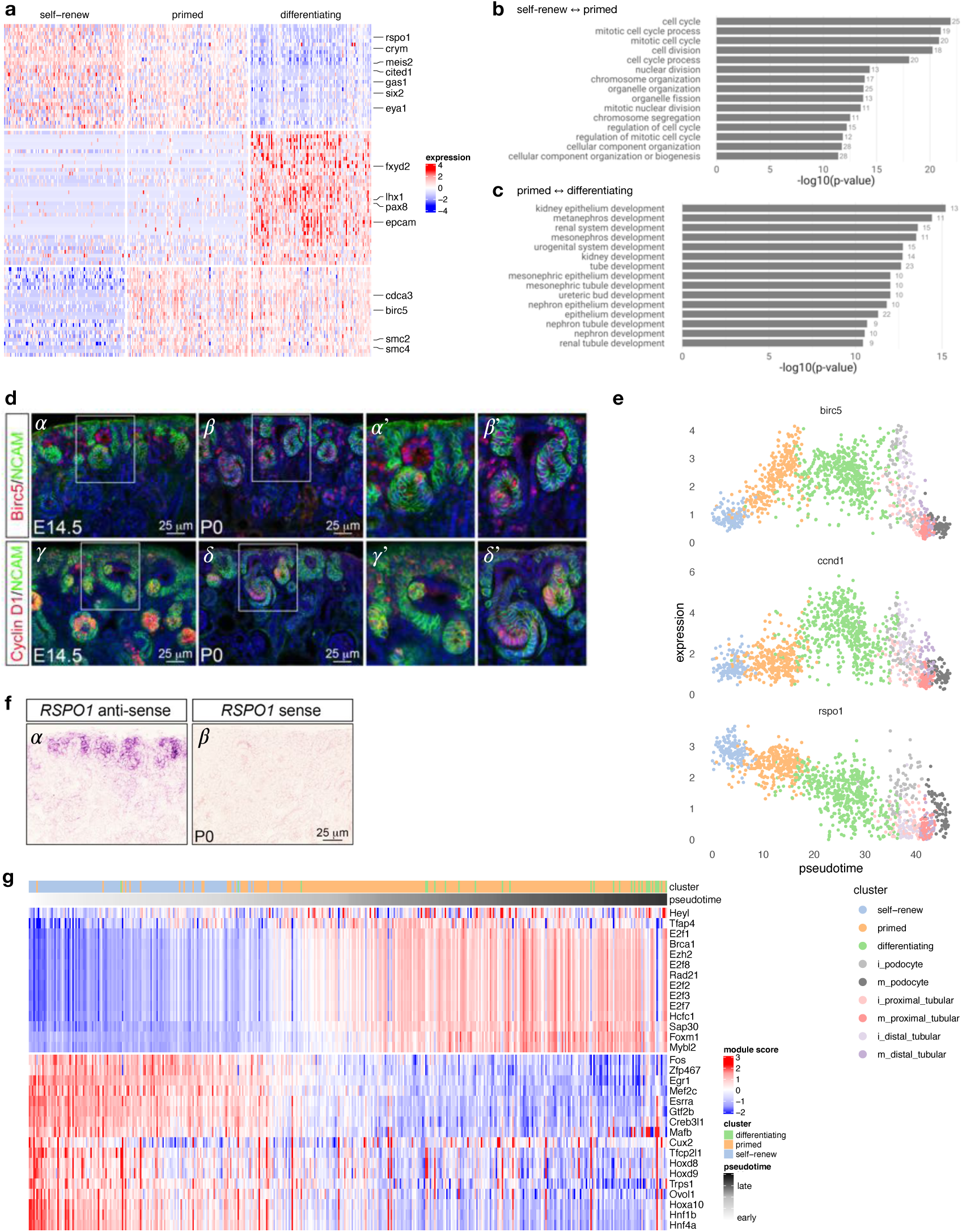
Transcriptional signatures of self-renewing, primed and differentiating nephron progenitor cells. Panel ***(a)*** shows differentially expressed genes on a heatmap of 100 random cells for each of the “self-renew”, “primed” and differentiating clusters, with key genes annotated on the right. Panels ***(b)*** and ***(C)*** show the 20 most-enriched Gene Ontology terms for genes differentially expressed between self-renewing and primed NP cells, and between primed NP cells and differentiating cells, respectively. ***(d)*** Immunofluorescence on kidney sections from embryonic day 14.5 (E14.5) and postnatal day 0 (P0) mice using anti-Birc5 *(α* - *β*′) and anti-Cyclin D1 *(γ - δ* ‘) antibodies (red). Nephron progenitors and their early epithelial derivatives were detected using an antibody against anti-Neural cell adhesion molecule (NCAM; green). Nuclei were counterstained with DAPI (blue). Scale bar, 25 μm. The sub-panels *α*′, *β*′, *γ*′ and *δ*’ are close-ups of the areas indicated by the white boxes. ***(e)*** Expression of *Birc5, Ccnd1* and *Rspo1* across pseudotime; colors indicate cell clusters. ***(f)*** *In situ* hybridization on cryosections of P0 kidneys confirms the expression of *Rspo1* in nephron progenitors and their early epithelial derivatives (*α*). No signal was detected with sense probe hybridization (*β*). Images are representative of three independent experiments. Scale bar 25 μm. ***(g)*** Inferred regulatory module activity based on SCENIC [49] across pseudotime for predominantly self-renewing and primed nephron progenitor cells.

### Transcriptional dynamics across nephron progenitor cell differentiation

#### Cell cycle activity distinguishes between two different types of nephron progenitor cells

Comparing gene expression between “self-renewing” with “primed” NP cells yielded cell cycle as a main difference between the two types of nephron progenitor cells (**Figure 3, Supplemental Table 3**). We find that cell cycle-related genes like *Birc5, Cdca3, Smc2* and *Smc4* are up-regulated between “primed” and “self-renewing” nephron progenitor cells. Immunofluorescence analysis on kidney sections from E14.5 and P0 mice indeed corroborates these observations, showing increased expression of Birc5 in pre-tubular aggregates/renal vesicles but negligible or absent expression in the “self-renewing” nephron progenitor cells (**Figure 3d** sub-panels *α, α*′, *β, β*′ and panel **3e**). These results corroborate previous findings that demonstrated that the committed nephron progenitor cells are more proliferative (=fast-cycling population) and more likely to differentiate than the slow-cycling, self-renewing NP population [45]. Next, comparing primed nephron progenitor cells with mixed/differentiating cells, we observe up-regulation of transcription factors associated with differentiation (*Lhx1, Pax8*), and down-regulation of nephron progenitor-associated genes like *Cited1, Six2, Eya1, Crym, Meis2, Rspo1* and others **(Figures 2b, 3a, 3e, Supplemental Table 4)**. *In situ* hybridization analysis confirms the expression of *Rspo1* [46] primarily in nephron progenitors (**Figure 3f**). Gene Ontology enrichment analysis comparing self-renewing with primed nephron progenitors, and primed progenitors with differentiating cells highlights differences in cell cycle-related biological processes between the two types of nephron progenitor cells and processes associated with differentiation between primed nephron progenitors and differentiating cells (**Figure 3b, c** and **Supplemental Table 5**).

To better understand these transcriptional changes occurring between self-renewing and primed nephron progenitor cells, we performed two additional types of gene set enrichment analyses utilizing lineage pseudotime annotation. Focusing on genes with changes across pseudotime (FWER < 0.01) we analyzed up-regulated and down-regulated genes separately. Performing enrichment analysis across Gene Ontology and Hallmark gene sets from MSigDB [47, 48] yielded “EPITHELIAL_MESENCHYMAL_TRANSITION” (FDR-adjusted p-value: 6.2E-5) as the most enriched hallmark gene set for the down-regulated genes, while gene sets enriched for up-regulated genes included “E2F_TARGETS” as the most enriched term as well as a multitude of gene sets associated with cell cycle/replication (**see Supplemental Table 6**). We then used SCENIC [49] to gain some insight into gene regulation driving the transcriptional changes we observe across pseudotime between the two nephron progenitor cell types. **Figure 3g** depicts the activity of inferred regulatory modules across pseudotime for 31 recovered transcription factors. We observe three regulatory modules of down-regulated genes, attributed to the transcription factors *Egr1, Maf* and *Fos*. These findings corroborate previous studies showing that these transcription factors are critical regulators of gene expression, controlling transition from a pluripotent to differentiated state in nephron progenitor and human embryonic stem cells [50]. For up-regulated genes, we observe modules associated with cell cycle-related transcription factors like *E2f8, Hcf1, Ezh2* and *Mybl2*, which have previously been implicated in specific aspects of cell cycle progression and cell fate decision in stem and progenitor cells [51-55]. Genes making up each module are provided in **Supplemental Table 8**.

#### *Birc5* expression in the tubular interconnection zone

The cell cycle-related genes *Birc5, Ccnd1* and *Tuba1a* were up-regulated in immature distal tubules (**Figures 2b, 4a**). Immunostaining analysis confirmed augmented expression of Birc5 and Cyclin D1 in the distal renal vesicle domain (**Figure 3d**). Interestingly, increased Birc5 was also observed in a sub-set of ureteric bud cells (**Supplemental Figure 4** and **Figure 3d**) located in the region of interconnection between the late renal vesicle and the adjacent ureteric bud tips. The fusion between the nephron and the collecting system is required for the formation of a functional renal network. Studies in mouse models have demonstrated that this process is driven by preferential cell division within the distal renal vesicle domain [56]. Therefore, Birc5 may contribute to tubular interconnection by regulating proliferation in the late renal vesicle and cell survival in the adjacent ureteric tip cells.

**Figure 4:**
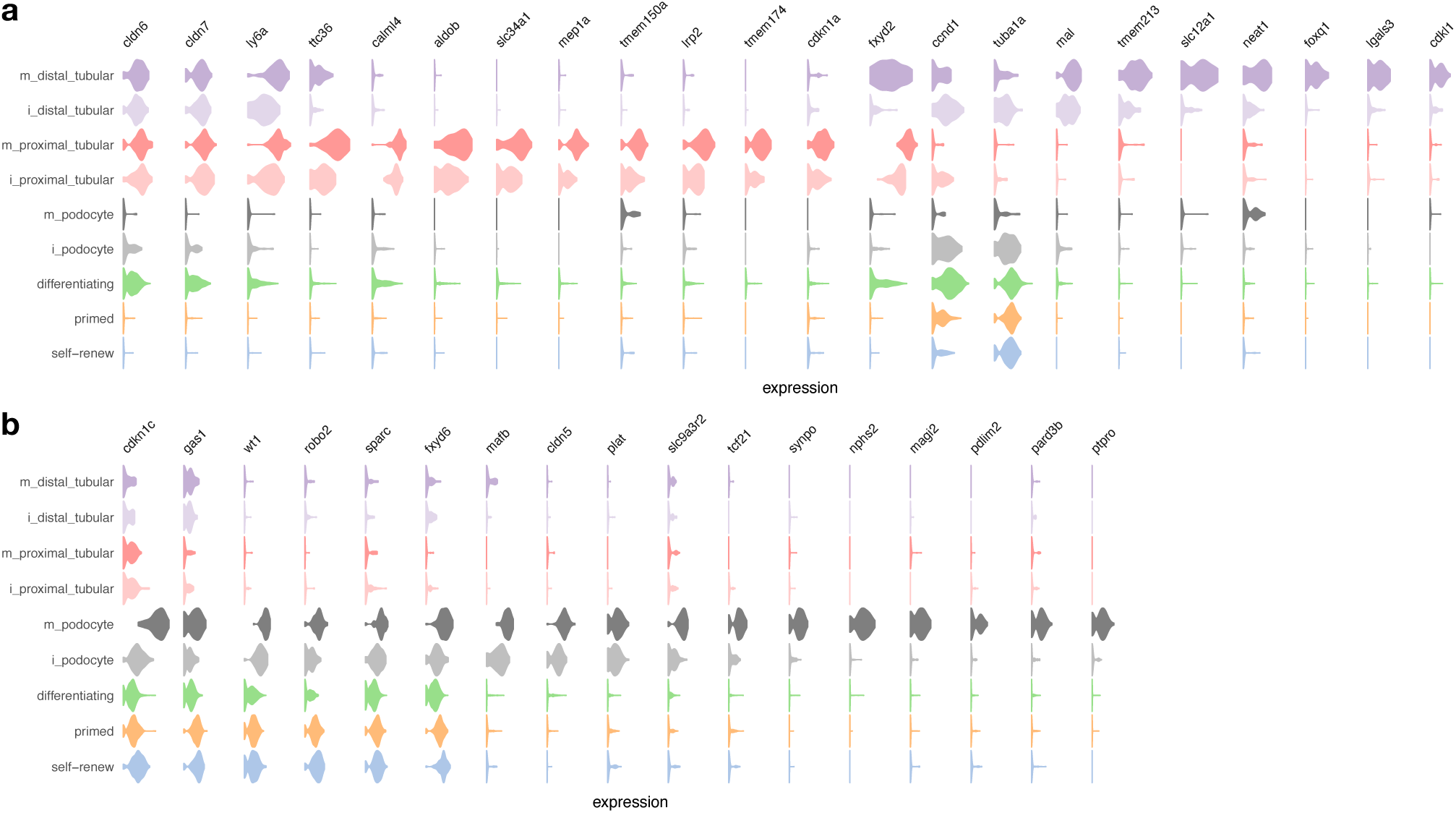
Transcriptional signatures of podocytes and tubular cells. ***(a)*** Violin plot of genes expressed in podocytes (rows are clusters and columns denote genes). ***(b)*** Same as *(a)*, but for proximal and distal tubular cells.

#### Conserved features in mouse and human podocyte development

In the podocyte lineage, the genes most significantly defining the cluster are *Pax8, Podxl* and *Nphs1. Podxl* and *Nphs1* (in combination with *Synpo, Nphs2* and *VEGF-A*) are restricted to a sub-population of mature podocytes [28, 31, 57], which is consistent with our observations (**Figures 2b and 4b**). In a sub-population of early podocytes, *WT1* and *Mafb* expression has been reported to overlap with the immature marker *Pax8* [27, 31], and is expressed in parietal epithelial cells [58], also consistent with our findings (**Figures 2b and 4b)**. For a summary of gene expression changes during podocyte development see **Supplemental Figure 5a** and **Supplemental Table 9**. Similar to previous scRNA-seq analysis in human fetal kidney [31], we observe enrichment in the PDZ domain proteins *Magi2, Slc9a3r2* and *Pard3b* in mature podocytes **(Figure 4b)**. We also observe podocyte-specific activity of *Cldn5* (while the claudins *Cldn6* and *Cldn7* are expressed in tubular lineages, **Figure 4a**). Further on, the gene *Sparc* (a cystine-rich matrix-associated protein) and the Tissue-Type Plasminogen Activator *Plat* are expressed specifically in the podocyte lineage (as is *Robo2*, a gene known to be expressed and colocalized with nephrin on the basal surface of mouse podocytes [59], while the cell-cycle regulator *Gas1* (Growth Arrest Specific 1) is expressed in undifferentiated cells and mature podocytes, but less so in mature tubular cells **(Figure 4b)**. Together, these findings further define the gene expression profile of the podocyte lineage, and they suggest substantial conservation between mouse and human developing podocytes.

**Figure 5:**
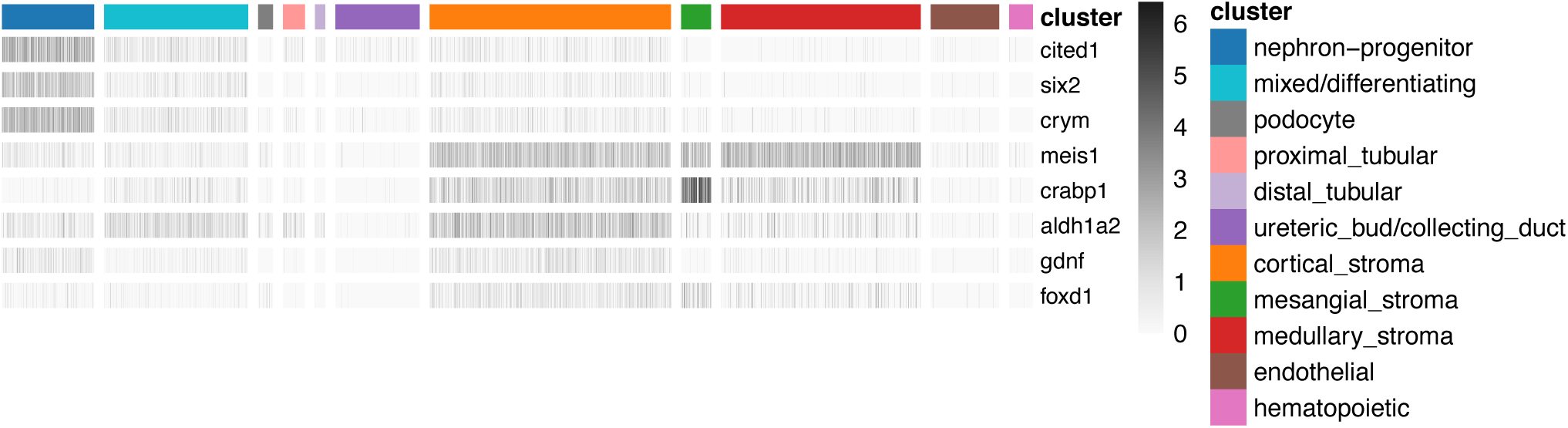
Expression of lineage-marker genes in unexpected cell types. Heatmap of gene expression (gray scale) of known lineage-marker genes (rows) across cells (columns), ordered by cell clusters (color index). We observe the expression of cap mesenchyme markers (*Cited1, Six2, Crym, Gdnf*) in stromal cells and vice versa, consistent with previous reports [27].

#### Gene expression differences between proximal and distal tubular cells

In addition to their respective marker genes *Pdzk1, Slc34a1, Tmem52b* and *Shd* (see above), we observe that tubular lineages express the claudins *Cldn6* and *Cldn7*, as well as the Lymphocyte Antigen 6 Complex Locus A (*Ly6a*, aka *Sca-1*), see **Figure 4a** (**Supplemental Figure 5b** and **Supplemental Table 10** show additional genes with differential expression between proximal and distal tubular cells). *Ly6a* is a member of the murine L6 family and has been reported to mark cancer and tissue-resident stem cells in mice [60]; however there is no known direct human ortholog for *Ly6a*, and also the function of the LU domain, which characterizes *Ly6a*’s superfamily of proteins, is currently unknown in humans [60].

We note that *Mep1a, Aldob* and *Tmem174* mark proximal tubular cells in our data (**Figure 4a**) and have been reported amongst the top-most down-regulated genes after *p53* conditional deletion in nephron progenitor cells [61]. Of the other three reported top down-regulated genes two (*Pck1* and *Cyp2d26*) also show proximal tubular cell specific expression (data not shown), while *Reg8* expression was not detected in our data. This is in line with the observation of fewer proximal tubular cells in P0 mutant kidneys reported in [61]. With respect to the cell cycle, we find that Cyclin-Dependent Kinase Inhibitor 1 A (*Cdkn1a*, aka *P21*) is active specifically in proximal tubular cells, while Cyclin-Dependent Kinase Inhibitor 1 C (*Cdnk1c*, aka *P57*) is primarily expressed in podocytes (**Figures 4a** and **4b**). For distal tubular cells, we don’t observe a selectively active kinase inhibitor but note that Cyclin-Dependent Kinase-like 1 (*Cdkl1*) is specifically expressed in this cell type. These findings pinpoint lineage-specific gene expression differences between the proximal vs. distal tubular lineages, and they point towards lineage-specific control of the cell cycle across nephron progenitor differentiation.

Recently published scRNA-seq papers have described differences in gene expression across a variety of proximal tubule transcripts and lncRNAs in different sexes in the adult mouse kidney [62, 63]. We observe that female-enriched markers, including *Gm4450, Lrp2, Sultd1, Aadat, Hao2* were highly expressed in our proximal tubular cluster, while most of the male-enriched markers (S*lc22a12, Cndp2*, C*esf1*, etc.) were absent or expressed at low levels. This data suggests that sexually dimorphic gene expression in proximal tubule may occur at or before E14.5.

### Expression of known lineage-marker genes in unexpected cell types

Expression of known lineage-marker genes in unexpected cell types has been reported based on the analysis of scRNA-seq data, for example that stromal cells express Gdnf [27]. Consistent with this report, we found that nephron progenitor markers (*Six2, Cited1, Crym*) are expressed in cells in the stromal cluster, and that stromal markers are present in the nephron progenitor cluster (*Meis1, Foxd1, Crabp1*). We also confirm that *Gdnf* is expressed in the stromal cluster (in addition to nephron progenitor cells), and that *Aldh1a2* RNA is present in stromal and nephron progenitor clusters **(Figure 5)**.

We note that nephron progenitor marker genes are not homogenously expressed across different stromal cell types. For instance, *Cited1* is detected (five or more reads) in about 7% of cortical stromal cells, but in less than 1% of other stromal cell types. We find similar enrichment (expression in ∼7% vs less than 1% of cells) for *Six2* and *Crym* in cortical stroma, whereas *Gdnf* is more modestly enriched in cortical stroma (expressed in ∼3% vs less than 1% of cells, respectively). We next focused on cortical stroma and looked at co-expression of nephron progenitor and stromal marker genes in the same cells (binary expression “on” vs. “off”) and find significant positive association between the expression of stromal-and nephron-progenitor genes (Fisher exact test, **Table 1**). This analysis demonstrates widespread co-expression of nephron-progenitor and stromal markers in the same cortical/stromal cells, and on average we observe higher odds ratios of association for *Col1a1* expression with nephron progenitor lineage-markers, compared with *Meis1* **(Table 1)**.

Further on, the cluster we identified as distal tubular cells contains cells with a distal-like expression profile, as characterized by the expression of *Tmem52b* and *Shd* [29]. However, despite the distinct lineage origins, cells from this cluster and from the ureteric_bud/collecting_duct cluster exhibit some transcriptional congruence [29, 64]. Specifically, *Calb1, Wdfc2* and *Mal* [28], which are thought to mark the ureteric but lineage, and *Mecom* [63], which is thought to mark distal tubular cells, are expressed in a significant fraction of cells in both these clusters, but absent in proximal tubular cells (**Table 2**).

## DISCUSSION

Over 30 terminally differentiated nephron cell types are required for the function of the mammalian kidney. The advent of scRNA-seq technology has made it possible to explore the cellular heterogeneity of the kidney and precisely identify the transcriptional signatures that define each of its cell types. In this study, we have performed scRNA-seq analysis of the developing mouse kidney at E14.5, a time point in which there is active nephron induction and varying degrees of nephron maturation. Major transcriptional clusters – corresponding to nephron progenitors, mixed/differentiating cells, podocytes, diferentiated tubules (proximal and distal), ureteric epithelium, stroma (medullary, mesangial and cortical), hematopoietic and endothelial lineages – are identified within the whole kidney analysis, and are consistent with prior single cell analyses of the developing mouse kidney [25-27, 29]. We find that cell cycle activity distinguishes between “primed” and “self-renewing” sub-populations of nephron progenitors. Furthermore, augmented *BirC5* expression occurs in immature distal tubules and a sub-set of ureteric bud cells, suggesting that *Birc5* might be a novel key molecule required for early events of tubular fusion between the distal nephron and the collecting duct epithelia.

All nephron segments derive from a multipotent self-renewing nephron progenitor population, which co-expresses the transcription factor *Six2* and the transcriptional activator *Cited1*. Previous studies have identified two sub-types of nephron progenitors, with Cited1^+^/ Six2^+^ progenitors transitioning to a Cited1^-^/ Six2^+^ primed state as the nephrogenesis proceeds [15, 34, 65]. Recent studies using time-lapse imaging and scRNA-seq analyses have indicated, however, that the nephron progenitor compartment is more heterogeneous that initially supposed [26, 27, 29, 66-68]. Moreover, differences in cell cycle length within progenitors appear to play a role in the sub-compartmentalization of the progenitor population [45]. In agreement, our scRNA-seq analysis shows separation of nephron progenitor cells into a “self-renewing” and “primed” sub-population, both co-expressing *Six2* and *Cited1*, but distinguished by higher cell cycle activity in the “primed” cells. Studies in mice have demonstrated that the committed nephron progenitors are more proliferative, exhibiting preferential exit from the cap mesenchyme compartment and differentiation into early nephrons [45]. Intriguingly, in the human renal cap mesenchyme, the “self-renewing” nephron progenitors exhibit a greater proliferative activity, compared to the committed progenitor population [32]. Although it is still unclear what drives these species-specific differences, this may be related to unique transcription factor expression in the human fetal kidney (such as continuous *Six1* expression in cap mesenchyme throughout nephrogenesis) [32].

We find that the transcriptional profile of “primed” nephron progenitors represents an intermediate/transitional state between self-renewing NP and mixed/differentiating cells (pre-tubular aggregates/renal vesicles), with lower levels of *Cited1* and increased expression of early commitment markers like *Lhx1* and markers of renal epithelia like *Pax8*. These findings are consistent with previous scRNA-seq analyses of developing human and mouse kidneys [28, 69], but are in contrast to other studies on nephron progenitor subpopulations where *Cited1* expression seems to be turned off prior to the activation of pre-tubular aggregate genes [15, 34, 68]. Such discrepancies might be due to differences in the technical sensitivity of the methods applied in each study (scRNA-seq versus immunofluorescence or *in situ* hybridization). They also highlight the importance of further analysis to confirm whether these nephron progenitor sub-populations coincide with distinct spatial domains within the developing kidney.

Our approach successfully identified a number of cell types in the developing kidney. Consistent with previously reported expression patterns, we observe *Podxl, Synpo, Nphs1* and *Nphs2* expression in mature podocytes, whereas *WT1* and *Mabf* are also expressed in a sub-population of early podocytes [31, 57]. Indeed, we also observed several PDZ domain proteins (*Magi2, Slc9a3r2* and *Pard3b*) [31] expressed in human developing podocytes in our data, suggesting that podocyte identity is conserved in the mouse and human developing kidney. Cells in the proximal tubular cluster are characterized by the specific expression of known proximal tubule markers, such as *Pdzk1* and *Slc34a1* [29, 63]. The scaffold protein Pdzk1 is essential for the proper localization of interacting proteins, such as the sodium-phosphate transporter NaPi-Iia (encoded by *Slc34a1*), in the brush border of the proximal tubular cells [70-72]. Interestingly, mutations in *Slc34a1* have been linked to nephrocalcinosis and Fanconi renotubular syndrome [73, 74]. Further on, we observe that *Cldn5* marks the podocyte lineage, while *Cldn6* and *Cldn7* are expressed in mixed/differentiating cells and both tubular lineages, but absent in podocytes. Genes specifically expressed in distal tubular cells in our data include Galectin 3 (*Lgals3*), *Slc12a1* and the long non-coding RNA *Neat1*.

The formation of a fully functional nephron entails fusion between the late renal vesicle and the adjacent ureteric tip. An elegant study using 3D modeling of nephrons and *Six2-eGFPCre* x *R26R-lacZ* mice demonstrated that this connecting segment of the nephron is derived from the cap mesenchyme (not the ureteric epithelium), and the process of fusion is likely driven by preferential cell division within the distal renal vesicle domain [56]. In line with this, our data identified augmented expression of cell cycle-related molecules, such as Cyclin D1 (*Ccnd1*), *Birc5* and *Tuba1a*, in immature distal tubules. Interestingly, high *Birc5* expression was also detected in ureteric bud cells located in the region where the ureteric tip connects with the distal portion of the renal vesicle (see **Supplemental Figure 4**).

*Birc5* (also known as Survivin) has been implicated in a number of kidney conditions, including autosomal-dominant polycystic kidney disease, acute kidney injury and renal cell carcinomas [75-79], however its role in context of normal kidney development is still unknown. In normal tissues, transcription of *Birc5* is tightly regulated in a cell cycle-dependent manner, reaching a peak in the G2/M phase [80-82], followed by a rapid decline at the G1 phase [83]. *Birc5* targets the chromosomal passenger complex (CPC) to the centromere, ultimately enabling proper chromosome segregation and cytokinesis [84-90]. *Birc5* also plays a role as an inhibitor of programmed cell death. Although this mechanism is not completely understood, it seems to require cooperation with other molecules (such as XIAP and HBXIP) and results in inhibition of caspase-9 [91-94]. Our data suggest that *Birc5* might be a novel key molecule required for early events of nephron patterning and fusion, by regulating cell survival and/or proliferation in late renal vesicle and the adjacent ureteric tip.

In line with other scRNA-seq studies, we identify three stromal clusters in our dataset: cortical, medullary and mesangial [25, 27]. The genes most significantly defining the mesangial stroma cluster are *Dlk1* and *Postn* [25, 29]. Cells in cortical stroma express high levels of *Aldh1a2* and *Dlk1*, while medullary stroma cluster contains cells with increased expression of *Cldn11*. The absence of an expression profile consistent with a loop of Henle signature in our scRNA-seq data is likely due to a low-abundance of these cell populations at E14.5 [95]. In addition, the lack of information on the cell diversity and identity within the loops of Henle continues to hinder the annotation of this segment [63].

In summary, this study provides an in-depth transcriptional profile of the developing mouse kidney at mid-gestation. Major main transcriptional clusters are identified, and are consistent with prior single cell analyses of the developing mouse kidney [25-27, 29]. Notably, we find that cell cycle activity distinguishes between the “primed” and “self-renewing” sub-populations of nephron progenitors, with increased levels of the cell cycle related genes *Birc5, Cdca3, Smc2* and *Smc4* in the “primed” sub-population. Finally, *BirC5* expression in immature distal tubules and ureteric bud cells may contribute to early events of tubular fusion between the distal nephron and the collecting duct epithelia.

## METHODS

### Embryonic kidney collection and single-cell RNA sequencing

Timed pregnant wild-type CD-1 female mice used in this study were obtained from Charles River Laboratories (Wilmington, MA, USA). The date on which the plug was observed was considered embryonic day 0.5 (E0.5). All experimental procedures were performed in accordance with the University of Pittsburgh Institutional Animal Care and Use Committee guidelines (IACUC protocol #17091432), which adheres to the NIH Guide for the Care and Use of Laboratory Animals.

We harvested two kidneys at E14.5 and generated a single cell suspension using 0.05% trypsin at 37°C for 10 minutes. Kidneys were mechanically dissociated with pipetting at 5 and 10 min. 3% fetal calf serum in PBS was added to halt the trypsin. The cell suspension was filtered using a 40 μm filter and pelleted. The cells were resuspended in 90% FCS in DMSO and frozen, prior to shipment to GENEWIZ Ing. Single-cell library preparation and sequencing was performed by GENEWIZ Inc. using the 10X Genomics Inc. Chromium 3’ Single Cell v2 library preparation kit. Cells exhibited high viability after freezing and thawing (>90%).

### Data processing, quality control, and normalization

#### Alignment and read counting

Sequencing data was processed using the cellranger count pipeline of the Cell Ranger software (version 2.2.0) (www.10xgenomics.com) to perform alignments and yield barcode and UMI counts, such that the cell detection algorithms are bypassed and counts for 10,000 cells are returned (force-cells=10000 option). The mouse reference genome (GRCm38.p4) and transcript annotations from Ensembl (version 84) were used [96].

#### Quality Control

The Bioconductor [97] R package DropletUtils [98, 99] was used to detect and remove empty droplets with default parameters at an FDR of 0.01, yielding a total of 5,887 non-empty droplets. Multiple quality control (QC) metrics were calculated using the R package scater [100] and cells with at least 1000 detected features, and percentage of mitochondrial counts less than 3 times the median absolute deviation (MAD) from the median value were considered, resulting in a total of 4,402 cells. We excluded putative doublets as the top 5% cells ranked by the hybrid score from the R package scds [101], further filtering out 220 droplets. Finally, only genes with at least three or more counts in at least three samples were considered, yielding a digital gene expression matrix comprising 11,155 genes in 4,183 cells/droplets.

#### Normalization

We normalized the data using size factors calculated using the deconvolution method implemented in the computeSumFactors function in the R package scran [102] after performing clustering using the quickcluster function on endogenous features with an average count >=0.1, (min.mean=0.1 option) yielding log-transformed normalized expression data. Feature selection and dimension reduction were performed using scran procedures. Briefly, we fit a mean-variance trend to the gene variances using the trendVar function and identified the biological component of the total variance with decomposeVar. All genes with an FDR < 0.01 and proportion of biological variance of at least 25% are considered as highly variable genes (HVG). Principal component analysis (PCA) was then performed using denoisePCA and two-dimensional representation was then derived using runTSNE.

### Identification of major structural components of the kidney

Cells were grouped into clusters using the scran R package by building a shared k-nearest-neighbors graph using buildSNNGraph (with use.dimred=PCA and k=25 options), followed by clustering with the Walktrap community finding algorithm as implemented in the igraph package (https://igraph.org), cutting the graph at 10 clusters. We used the expression of a curated list of marker genes for major components of the developing kidney (see **Figure 1c**) to assign cluster labels. Cluster-specific markers were derived using the find-Markers function (**Supplemental Table 1**). We note that at this resolution tubular distal cells were grouped in the mixed/differentiating group; specific analysis of nephron progenitor descendant cell types then revealed distinct groups of distal vs. proximal tubular cells (see below).

### Nephron progenitor and descendant cell types

#### Selecting and characterizing NP lineage cells

Focusing on nephron progenitor and descendant cell types (termed “nephron-progenitor”, “mixed/differentiating” (at that point containing “distal_tubular” cells as well), “podocytes and “proximal_tubular” in **Figure 1**) and requiring expression of each gene with at least three counts in three cells yielded a gene expression matrix of 9,611 genes across 1,273 cells. Following the same procedure as before we derived a low-dimensional representation and identified six clusters of cells, corresponding to two types of nephron progenitor cells (“self-renew” and “primed”), “mixed/differentiating” cells as well as distal tubular cells and proximal tubular cells (**Figure 2**). Cluster-specific marker genes were derived as before and are reported in **Supplemental Table 2**. Enrichment analysis for Gene Ontology terms enriched between “self-renew” and “primed” and between “primed” and “mixed/differentiating” (**Figures 3b and 3c, Supplemental Tables 3 - 5**) were performed using the topGO function of the limma Bioconductor package [103] with default parameters. We also used SAVER [104] to impute gene expression values across this set of cells, which we then utilized in pseudotime-related analyses described below.

#### Pseudotime Analysis of NP cells

Pseudotime analysis was performed using slingshot [105], using cluster labels and principal components derived as described above (via the clusterLabels and reducedDim options). This recovered three lineages (to podocytes, distal-, and proximal tubular cells), with cells in “self-renew”, “primed” and early “differentiating/mixed” being shared (see **Supplemental Figure 2**).

Next, we fitted a multinomial log-linear model (using the nnet package [106]) relating pseudotime with the annotated clusters. For cells with more than one annotated lineage (in the “self-renew”, “primed” and early “differentiating” clusters) lineage-pseudotimes from slingshot were averaged. This enabled us to define *NP-cells* as cells with annotated pseudotime less than the (pseudo)timepoint between “primed” and “differentiating” where the probability of the “primed” cluster has declined to 50% (i.e., 50% probability for “differentiating”. These cells contain all “self-renew” cells, 28 “differentiating” cells and all but 15 “primed” cells and were used in subsequent pseudotime analyses comparing self-renewing and differentiating cells.

We then applied generalized additive models, as implemented in the mgcv package [107], to screen for pseudotime-associated genes (**Figure 3g**) by modeling gene expression as a smooth function of pseudotime. Focusing on high-quality pseudotime-associated genes (FWER<1% modeling significance, plus highly-expressed and with an absolute spearman correlation of gene expression with pseudotime larger than 0.4) yielded 168 genes with overall decreasing expression across pseudotime (down-regulated), and 399 with increasing expression (up-regulated). Gene sets are reported in **Supplemental Table 6**. We then used MsigDB (v7.0) [47] and hypergeometric tests to screen for annotated gene sets enriched for up- or down-regulated genes, focusing on Gene Ontology and Hallmark gene sets (**Supplemental Table 7**). Finally we screened for regulatory modules in time-varying genes using SCENIC [49], where we used default options including GENIE3 for [108] network inference (see **Figure 3**, and **Supplemental Table 8** lists the modules we recovered).

#### Immunohistochemical staining

Kidneys dissected from embryonic day 14.5 (E14.5) and postnatal day 0 (P0) mice were fixed overnight in 4% paraformaldehyde, embedded in paraffin and sectioned at 4 μm. After deparaffinization, rehydration, and permeabilization in PBS-Tween (PBS-T), antigen retrieval was performed by boiling the slides in 10 mM sodium citrate pH 6.0 buffer for 30 min. Next, sections were blocked in 3% bovine serum albumin (BSA) and incubated overnight with antibodies recognizing Birc5 (#2808, Cell Signaling Technology, Danvers, MA, USA), Cyclin D1 (#2978, Cell Signaling) and Neural cell adhesion molecule (C9672, Sigma-Aldrich, St. Louis, MO, USA) at the dilutions recommended by the manufactures. On the next day, sections were washed with PBS-T, incubated with secondary antibodies at the dilution of 1:200, washed again with PBS-T, and mounted in Fluoro Gel with DABCO (Electron Microscopy Science, Hatfield, PA) before being visualized with a Leica DM2500 microscope and photographed with a Leica DFC 7000T camera using LAS X software (Leica, Buffalo Grove, IL, USA). Goat anti-rabbit 594 (#111-515-047) and donkey anti-mouse 488 (#715-545-151) antibodies were purchased from Jackson InmmunoResearch Laboratories (West Grove, PA, USA).

#### *In situ* hybridization

Kidneys were harvested from P0 pups, fixed in 4% paraformaldehyde overnight, treated with 30% sucrose/PBS and embedded in Tissue-Tek Optimal Cutting Temperature Compound (OCT; Sakura, Torrance, CA, USA). *In situ* hybridization was conducted on 10 μm cryosections as described [109]. To generate sense and antisense probes, plasmids were linearized and transcribed as follows: *pGEM*^***®***^*-T Easy-RSPO1-SacII/SP6 and pGEM*^***®***^*-T Easy-RSPO1-SaI/T7*.

### Reproducibility and data availability

Computer code used for data processing and data analysis is available on github (https://github.com/kostkalab/wksc_manuscript). Single cell RNA sequencing data is available on GEO (GEO identifier to be determined)

## ACKNOWLEDGEMENTS

This work was supported by NIH grants to D Kostka (NIGMS GM115836) and J. Ho (NIDDK DK103776). D. M. Cerqueira was supported by Nephrotic Syndrome Study Network (NEPTUNE) Career Development Award and Children’s Hospital of Pittsburgh Research Advisory Council Postdoctoral Fellowship. A. Clugston was supported by NIDDK T32 (DK061296) Institutional National Research Service Award.

## AUTHOR CONTRIBUTIONS

A. Bais and A. Clugston designed experiments and analyzed data; D. M. Cerqueira performed experiments and revised the paper; J. Ho and D. Kostka designed experiments, analyzed the data and wrote the paper. All authors reviewed, revised, and approved the paper prior to submission.

## ADDITIONAL INFORMATION

The authors declare no competing interests.

## SUPPLEMENTAL MATERIAL

**Supplemental Figure 1: *Heatmap of marker genes that distinguish between kidney cell types***. Rows are genes (fifteen top-most marker genes have been selected for each cluster), and columns are cells grouped by cell types.

**Supplemental Figure 2: *Differentiation lineages for nephron progenitor cells***. A tSNE embedding for nephron progenitor cells is shown. Differentiation lineages inferred by the slingshot R package are displayed in gray.

**Supplemental Figure 3: *Differentially expressed genes during nephron progenitor cell differentiation***. Heatmap of top differentially expressed genes between annotated clusters (column labels).

**Supplemental Figure 4: *Consistent* Birc5 *expression in distal tubular cells and a subpopulation of cells from the ureteric bud***. Shown are tSNE plots of nephron progenitor derived cells and cells of the ureteric bud / collecting duct (ub/cd) cluster (see **Figure 1**). ***(a)*** Cell-type annotations for depicted cells. ***(b)*** *Birc5* expression. Arrows denote early distal tubular cells and ub/cd cells with similar *Birc5* expression.

**Supplemental Figure 5: *Differentially expressed genes in the podocyte and tubular lineages. (a)*** Heatmap of top 100 differentially expressed genes between immature and mature podocytes and ***(b)*** between proximal and distal tubular cells. ***(c)*** Differential gene expression between immature and mature proximal tubular cells, and ***(d)*** between immature and mature distal tubular cells.

**Supplemental Table 1: *Marker genes that distinguish between kidney cell types***.

For each cluster marker genes with an FDR < 0.05 are reported (see **Figure 1**).

**Supplemental Table 2: *Marker genes that distinguish nephron-progenitor derived cell types***. For each cluster (imperfect) marker genes with an FDR < 0.05 are reported (see **Figure 2**).

**Supplemental Table 3: *Differentially expressed genes for self-renewing vs. primed nephron progenitor cell types***. For each comparison, differentially expressed genes (FDR < 0.1) are reported (see **Figure 3a**).

**Supplemental Table 4: *Differentially expressed genes for primed nephron progenitor cells vs. differentiating cells***. Differentially expressed genes (FDR < 0.1) are reported (see **Figure 3a**).

**Supplemental Table 5: *Gene Ontology terms enriched for differentially expressed genes, comparing self-renewing vs. primed nephron progenitor cells and primed vs. differentiating cells***. See **Figure 3b and 3c**.

**Supplemental Table 6: *Genes associated with pseudotime across nephron progenitor cells***. Genes associated with pseudotime are reported (FWER < 1%).

**Supplemental Table 7: *Gene sets enriched for genes increasing with pseudotime and for genes decreasing with pseudotime across nephron progenitor cells***. Gene sets from MSigDB for Gene Ontology (Biological Process) and Hallmark gene sets (FDR < 0.01) are reported.

**Supplemental Table 8: *Regulatory modules***. Regulatory modules of genes active in the transition between self-renewing and primed nephron progenitor cells (see **Figure 3g**).

**Supplemental Table 9: *Differentially expressed genes for mature vs. immature nephron progenitor derived cell types***. Differentially expressed genes (FDR < 0.1) are reported (see **Supplemental Figure 5**).

**Supplemental Table 10: *Differentially expressed genes for distal vs. proximal tubular cells***. Differentially expressed genes (FDR < 0.1) are reported (see **Supplemental Figure 5**).

## REFERENCES

1. Capone, V.P., et al., Genetics of Congenital Anomalies of the Kidney and Urinary Tract: The Current State of Play. International Journal of Molecular Sciences, 2017.18(4): p. 796.

2. Yosypiv, I.V., Congenital anomalies of the kidney and urinary tract: a genetic disorder? International journal of nephrology, 2012. 2012: p. 909083.

3. Keller, G., et al., Nephron number in patients with primary hypertension. N Engl J Med, 2003.348(2): p. 101–8.

4. Hoy, W.E., et al., Reduced nephron number and glomerulomegaly in Australian Aborigines: a group at high risk for renal disease and hypertension. Kidney Int, 2006.70(1): p. 104–10.

5. Humphreys, B.D., Mechanisms of Renal Fibrosis. Annu Rev Physiol, 2018. 80: p. 309–326.

6. Coresh, J., et al., Prevalence of chronic kidney disease in the United States. JAMA, 2007.298(17): p. 2038–47.

7. Boyle, S., et al., Fate mapping using Cited1-CreERT2 mice demonstrates that the cap mesenchyme contains self-renewing progenitor cells and gives rise exclusively to nephronic epithelia. Dev Biol, 2008.313(1): p. 234–45.

8. Kobayashi, A., et al., Six2 defines and regulates a multipotent self-renewing nephron progenitor population throughout mammalian kidney development. Cell Stem Cell, 2008.3(2): p. 169–81.

9. Little, M.H. and A.P. McMahon, Mammalian kidney development: principles, progress, and projections. Cold Spring Harb Perspect Biol, 2012. 4(5).

10. Brown, A.C., et al., FGF/EGF signaling regulates the renewal of early nephron progenitors during embryonic development. Development, 2011.138(23): p. 5099–112.

11. Barak, H., et al., FGF9 and FGF20 maintain the stemness of nephron progenitors in mice and man. Dev Cell, 2012.22(6): p. 1191–207.

12. Muthukrishnan, S.D., et al., Concurrent BMP7 and FGF9 signalling governs AP-1 function to promote self-renewal of nephron progenitor cells. Nat Commun, 2015. 6: p. 10027.

13. Di Giovanni, V., et al., Fibroblast growth factor receptor-Frs2alpha signaling is critical for nephron progenitors. Dev Biol, 2015.400(1): p. 82–93.

14. Blank, U., et al., BMP7 promotes proliferation of nephron progenitor cells via a JNK-dependent mechanism. Development, 2009.136(21): p. 3557–66.

15. Brown, A.C., et al., Role for compartmentalization in nephron progenitor differentiation. Proc Natl Acad Sci U S A, 2013.110(12): p. 4640–5.

16. Karner, C.M., et al., Canonical Wnt9b signaling balances progenitor cell expansion and differentiation during kidney development. Development, 2011.138(7): p. 1247–57.

17. Park, J.S., et al., Six2 and Wnt regulate self-renewal and commitment of nephron progenitors through shared gene regulatory networks. Dev Cell, 2012.23(3): p. 637–51.

18. Majumdar, A., et al., Wnt11 and Ret/Gdnf pathways cooperate in regulating ureteric branching during metanephric kidney development. Development, 2003.130(14): p. 3175–85.

19. Dressler, G.R., The cellular basis of kidney development. Annu Rev Cell Dev Biol, 2006. 22: p. 509–29.

20. Schell, C., N. Wanner, and T.B. Huber, Glomerular development--shaping the multi-cellular filtration unit. Semin Cell Dev Biol, 2014. 36: p. 39–49.

21. Lindahl, P., et al., Paracrine PDGF-B/PDGF-Rbeta signaling controls mesangial cell development in kidney glomeruli. Development, 1998.125(17): p. 3313–22.

22. Betsholtz, C., et al., Role of platelet-derived growth factor in mesangium development and vasculopathies: lessons from platelet-derived growth factor and platelet-derived growth factor receptor mutations in mice. Curr Opin Nephrol Hypertens, 2004.13(1): p. 45–52.

23. Robert, B., X. Zhao, and D.R. Abrahamson, Coexpression of neuropilin-1, Flk1, and VEGF(164) in developing and mature mouse kidney glomeruli. Am J Physiol Renal Physiol, 2000.279(2): p. F275–82.

24. Abrahamson, D.R., Glomerulogenesis in the developing kidney. Semin Nephrol, 1991.11(4): p. 375–89.

25. Adam, M., A.S. Potter, and S.S. Potter, Psychrophilic proteases dramatically reduce single-cell RNA-seq artifacts: a molecular atlas of kidney development. Development, 2017.144(19): p. 3625–3632.

26. Brunskill, E.W., et al., Single cell dissection of early kidney development: multilineage priming. Development, 2014.141(15): p. 3093–101.

27. Magella, B., et al., Cross-platform single cell analysis of kidney development shows stromal cells express Gdnf. Dev Biol, 2018.434(1): p. 36–47.

28. Combes, A.N., et al., Single cell analysis of the developing mouse kidney provides deeper insight into marker gene expression and ligand-receptor crosstalk. Development, 2019. 146(12).

29. Combes, A.N., et al., Correction: Single cell analysis of the developing mouse kidney provides deeper insight into marker gene expression and ligand-receptor crosstalk (doi:10.1242/dev.178673). Development, 2019. 146(13).

30. England, A.R., et al., Identification and characterization of cellular heterogeneity within the developing renal interstitium. Development, 2020. 147(15).

31. Menon, R., et al., Single-cell analysis of progenitor cell dynamics and lineage specification in the human fetal kidney. Development, 2018. 145(16).

32. Wang, P., et al., Dissecting the Global Dynamic Molecular Profiles of Human Fetal Kidney Development by Single-Cell RNA Sequencing. Cell Rep, 2018.24(13): p. 3554–3567 e3.

33. Lindstrom, N.O., et al., Conserved and Divergent Features of Mesenchymal Progenitor Cell Types within the Cortical Nephrogenic Niche of the Human and Mouse Kidney. J Am Soc Nephrol, 2018.

34. Brown, A.C., S.D. Muthukrishnan, and L. Oxburgh, A synthetic niche for nephron progenitor cells. Dev Cell, 2015.34(2): p. 229–41.

35. Tanigawa, S., et al., Selective In Vitro Propagation of Nephron Progenitors Derived from Embryos and Pluripotent Stem Cells. Cell Rep, 2016.15(4): p. 801–813.

36. Li, Z., et al., 3D Culture Supports Long-Term Expansion of Mouse and Human Nephrogenic Progenitors. Cell Stem Cell, 2016.19(4): p. 516–529.

37. Takasato, M. and M.H. Little, A strategy for generating kidney organoids: Recapitulating the development in human pluripotent stem cells. Dev Biol, 2016.420(2): p. 210–220.

38. Morizane, R. and J.V. Bonventre, Generation of nephron progenitor cells and kidney organoids from human pluripotent stem cells. Nat Protoc, 2017.12(1): p. 195–207.

39. Hartman, H.A., H.L. Lai, and L.T. Patterson, Cessation of renal morphogenesis in mice. Dev Biol, 2007.310(2): p. 379–87.

40. Marlier, A. and T. Gilbert, Expression of retinoic acid-synthesizing and - metabolizing enzymes during nephrogenesis in the rat. Gene Expr Patterns, 2004.5(2): p. 179–85.

41. Harding, S.D., et al., The GUDMAP database--an online resource for genitourinary research. Development, 2011.138(13): p. 2845–53.

42. Diez-Roux, G., et al., A high-resolution anatomical atlas of the transcriptome in the mouse embryo. PLoS Biol, 2011.9(1): p. e1000582.

43. Takemoto, M., et al., Large-scale identification of genes implicated in kidney glomerulus development and function. EMBO J, 2006.25(5): p. 1160–74.

44. Lee, J.W., C.L. Chou, and M.A. Knepper, Deep Sequencing in Microdissected Renal Tubules Identifies Nephron Segment-Specific Transcriptomes. J Am Soc Nephrol, 2015.26(11): p. 2669–77.

45. Short, K.M., et al., Global quantification of tissue dynamics in the developing mouse kidney. Dev Cell, 2014.29(2): p. 188–202.

46. Lindstrom, N.O., et al., Conserved and Divergent Molecular and Anatomic Features of Human and Mouse Nephron Patterning. J Am Soc Nephrol, 2018.29(3): p. 825–840.

47. Liberzon, A., et al., The Molecular Signatures Database (MSigDB) hallmark gene set collection. Cell Syst, 2015.1(6): p. 417–425.

48. Liberzon, A., et al., Molecular signatures database (MSigDB) 3.0. Bioinformatics, 2011.27(12): p. 1739–40.

49. Aibar, S., et al., SCENIC: single-cell regulatory network inference and clustering. Nat Methods, 2017.14(11): p. 1083–1086.

50. Chen, P.M., et al., c-Maf regulates pluripotency genes, proliferation/self-renewal, and lineage commitment in ROS-mediated senescence of human mesenchymal stem cells. Oncotarget, 2015.6(34): p. 35404–18.

51. Julian, L.M. and A. Blais, Transcriptional control of stem cell fate by E2Fs and pocket proteins. Front Genet, 2015. 6: p. 161.

52. Hu, T., et al., Concomitant inactivation of Rb and E2f8 in hematopoietic stem cells synergizes to induce severe anemia. Blood, 2012.119(19): p. 4532–42.

53. Jolly, L.A., et al., HCFC1 loss-of-function mutations disrupt neuronal and neural progenitor cells of the developing brain. Hum Mol Genet, 2015.24(12): p. 3335–47.

54. Chen, Y.H., M.C. Hung, and L.Y. Li, EZH2: a pivotal regulator in controlling cell differentiation. Am J Transl Res, 2012.4(4): p. 364–75.

55. Papetti, M. and L.H. Augenlicht, MYBL2, a link between proliferation and differentiation in maturing colon epithelial cells. J Cell Physiol, 2011.226(3): p. 785–91.

56. Georgas, K., et al., Analysis of early nephron patterning reveals a role for distal RV proliferation in fusion to the ureteric tip via a cap mesenchyme-derived connecting segment. Dev Biol, 2009.332(2): p. 273–86.

57. Bariety, J., et al., Parietal podocytes in normal human glomeruli. J Am Soc Nephrol, 2006.17(10): p. 2770–80.

58. Ohse, T., et al., De novo expression of podocyte proteins in parietal epithelial cells during experimental glomerular disease. Am J Physiol Renal Physiol, 2010.298(3): p. F702–11.

59. Fan, X., et al., Inhibitory effects of Robo2 on nephrin: a crosstalk between positive and negative signals regulating podocyte structure. Cell Rep, 2012.2(1): p. 52–61.

60. Upadhyay, G., Emerging Role of Lymphocyte Antigen-6 Family of Genes in Cancer and Immune Cells. Front Immunol, 2019. 10: p. 819.

61. Li, Y., et al., p53 Enables metabolic fitness and self-renewal of nephron progenitor cells. Development, 2015.142(7): p. 1228–41.

62. Wu, H., et al., Proximal Tubule Translational Profiling during Kidney Fibrosis Reveals Proinflammatory and Long Noncoding RNA Expression Patterns with Sexual Dimorphism. J Am Soc Nephrol, 2020.31(1): p. 23–38.

63. Ransick, A., et al., Single-Cell Profiling Reveals Sex, Lineage, and Regional Diversity in the Mouse Kidney. Dev Cell, 2019.51(3): p. 399–413 e7.

64. Combes, A.N., et al., Single-cell analysis reveals congruence between kidney organoids and human fetal kidney. Genome Med, 2019.11(1): p. 3.

65. O’Brien, L.L., et al., Wnt11 directs nephron progenitor polarity and motile behavior ultimately determining nephron endowment. Elife, 2018. 7.

66. Combes, A.N., et al., Cap mesenchyme cell swarming during kidney development is influenced by attraction, repulsion, and adhesion to the ureteric tip. Dev Biol, 2016.418(2): p. 297–306.

67. Lawlor, K.T., et al., Nephron progenitor commitment is a stochastic process influenced by cell migration. Elife, 2019. 8.

68. Mugford, J.W., et al., High-resolution gene expression analysis of the developing mouse kidney defines novel cellular compartments within the nephron progenitor population. Dev Biol, 2009.333(2): p. 312–23.

69. Lindstrom, N.O., et al., Conserved and Divergent Features of Mesenchymal Progenitor Cell Types within the Cortical Nephrogenic Niche of the Human and Mouse Kidney. J Am Soc Nephrol, 2018.29(3): p. 806–824.

70. Thomson, R.B., et al., Role of PDZK1 in membrane expression of renal brush border ion exchangers. Proc Natl Acad Sci U S A, 2005.102(37): p. 13331–6.

71. Anzai, N., et al., The multivalent PDZ domain-containing protein PDZK1 regulates transport activity of renal urate-anion exchanger URAT1 via its C terminus. J Biol Chem, 2004.279(44): p. 45942–50.

72. Sayer, J.A., Progress in Understanding the Genetics of Calcium-Containing Nephrolithiasis. J Am Soc Nephrol, 2017.28(3): p. 748–759.

73. Fearn, A., et al., Clinical, biochemical, and pathophysiological analysis of SLC34A1 mutations. Physiol Rep, 2018.6(12): p. e13715.

74. Chau, H., et al., Renal calcification in mice homozygous for the disrupted type IIa Na/Pi cotransporter gene Npt2. J Bone Miner Res, 2003.18(4): p. 644–57.

75. AbouAlaiwi, W.A., et al., Endothelial cells from humans and mice with polycystic kidney disease are characterized by polyploidy and chromosome segregation defects through survivin down-regulation. Hum Mol Genet, 2011.20(2): p. 354–67.

76. Chen, J., et al., Survivin mediates renal proximal tubule recovery from AKI. J Am Soc Nephrol, 2013.24(12): p. 2023–33.

77. Zhou, D., et al., Tubule-specific ablation of endogenous beta-catenin aggravates acute kidney injury in mice. Kidney Int, 2012.82(5): p. 537–47.

78. Byun, S.S., et al., Expression of survivin in renal cell carcinomas: association with pathologic features and clinical outcome. Urology, 2007.69(1): p. 34–7.

79. Parker, A.S., et al., Comparison of digital image analysis versus visual assessment to assess survivin expression as an independent predictor of survival for patients with clear cell renal cell carcinoma. Hum Pathol, 2008.39(8): p. 1176–84.

80. Kobayashi, K., et al., Expression of a murine homologue of the inhibitor of apoptosis protein is related to cell proliferation. Proc Natl Acad Sci U S A, 1999.96(4): p. 1457–62.

81. Li, F. and D.C. Altieri, Transcriptional analysis of human survivin gene expression. Biochem J, 1999. 344 Pt 2: p. 305–11.

82. Li, F., et al., Control of apoptosis and mitotic spindle checkpoint by survivin. Nature, 1998.396(6711): p. 580–4.

83. Zhao, J., et al., The ubiquitin-proteasome pathway regulates survivin degradation in a cell cycle-dependent manner. J Cell Sci, 2000. 113 Pt 23: p. 4363–71.

84. Uren, A.G., et al., Survivin and the inner centromere protein INCENP show similar cell-cycle localization and gene knockout phenotype. Curr Biol, 2000.10(21): p. 1319–28.

85. Lens, S.M., et al., Survivin is required for a sustained spindle checkpoint arrest in response to lack of tension. EMBO J, 2003.22(12): p. 2934–47.

86. Carvalho, A., et al., Survivin is required for stable checkpoint activation in taxol-treated HeLa cells. J Cell Sci, 2003. 116(Pt 14): p. 2987–98.

87. Rajagopalan, S. and M.K. Balasubramanian, Schizosaccharomyces pombe Bir1p, a nuclear protein that localizes to kinetochores and the spindle midzone, is essential for chromosome condensation and spindle elongation during mitosis. Genetics, 2002.160(2): p. 445–56.

88. Yue, Z., et al., Deconstructing Survivin: comprehensive genetic analysis of Survivin function by conditional knockout in a vertebrate cell line. J Cell Biol, 2008.183(2): p. 279–96.

89. Speliotes, E.K., et al., The survivin-like C. elegans BIR-1 protein acts with the Aurora-like kinase AIR-2 to affect chromosomes and the spindle midzone. Mol Cell, 2000.6(2): p. 211–23.

90. Yang, D., A. Welm, and J.M. Bishop, Cell division and cell survival in the absence of survivin. Proc Natl Acad Sci U S A, 2004.101(42): p. 15100–5.

91. Marusawa, H., et al., HBXIP functions as a cofactor of survivin in apoptosis suppression. EMBO J, 2003.22(11): p. 2729–40.

92. Dohi, T., et al., An IAP-IAP complex inhibits apoptosis. J Biol Chem, 2004.279(33): p. 34087–90.

93. Mita, A.C., et al., Survivin: key regulator of mitosis and apoptosis and novel target for cancer therapeutics. Clin Cancer Res, 2008.14(16): p. 5000–5.

94. Wheatley, S.P. and D.C. Altieri, Survivin at a glance. J Cell Sci, 2019. 132(7).

95. Chang, C.H. and J.A. Davies, In developing mouse kidneys, orientation of loop of Henle growth is adaptive and guided by long-range cues from medullary collecting ducts. J Anat, 2019.235(2): p. 262–270.

96. Zerbino, D.R., et al., Ensembl 2018. Nucleic Acids Res, 2018. 46(D1): p. D754–D761.

97. Huber, W., et al., Orchestrating high-throughput genomic analysis with Bioconductor. Nat Methods, 2015.12(2): p. 115–21.

98. Lun, A.T.L., et al., EmptyDrops: distinguishing cells from empty droplets in droplet-based single-cell RNA sequencing data. Genome Biol, 2019.20(1): p. 63.

99. Griffiths, J.A., et al., Detection and removal of barcode swapping in single-cell RNA-seq data. Nat Commun, 2018.9(1): p. 2667.

100. McCarthy, D.J., et al., Scater: pre-processing, quality control, normalization and visualization of single-cell RNA-seq data in R. Bioinformatics, 2017.33(8): p. 1179–1186.

101. Bais, A.S. and D. Kostka, scds: computational annotation of doublets in single-cell RNA sequencing data. Bioinformatics, 2020.36(4): p. 1150–1158.

102. Lun, A.T., D.J. McCarthy, and J.C. Marioni, A step-by-step workflow for low-level analysis of single-cell RNA-seq data with Bioconductor. F1000Res, 2016. 5: p. 2122.

103. Ritchie, M.E., et al., limma powers differential expression analyses for RNA-sequencing and microarray studies. Nucleic Acids Res, 2015.43(7): p. e47.

104. Huang, M., et al., SAVER: gene expression recovery for single-cell RNA sequencing. Nat Methods, 2018.15(7): p. 539–542.

105. Street, K., et al., Slingshot: cell lineage and pseudotime inference for single-cell transcriptomics. BMC Genomics, 2018.19(1): p. 477.

106. Venables, W.N.R. B. D, Modern Applied Statistics with S. Fourth ed. 2002, New York: Springer.

107. B., W.S.N.P.N.S., Smoothing parameter and model selection for general smooth models (with discussion). Journal of the American Statistical Association, 2016. 111: p. 1548–1575.

108. Huynh-Thu, V.A., et al., Inferring regulatory networks from expression data using tree-based methods. PLoS One, 2010. 5(9).

109. D, W., In Situ Hybridization: A Practical Approach. 1992: IRL Press, Oxford University

