## Supplemental Figures for "Single cell RNA sequencing reveals differential cell cycle activity in key cell populations during nephrogenesis"

### Supplemental Figure 1

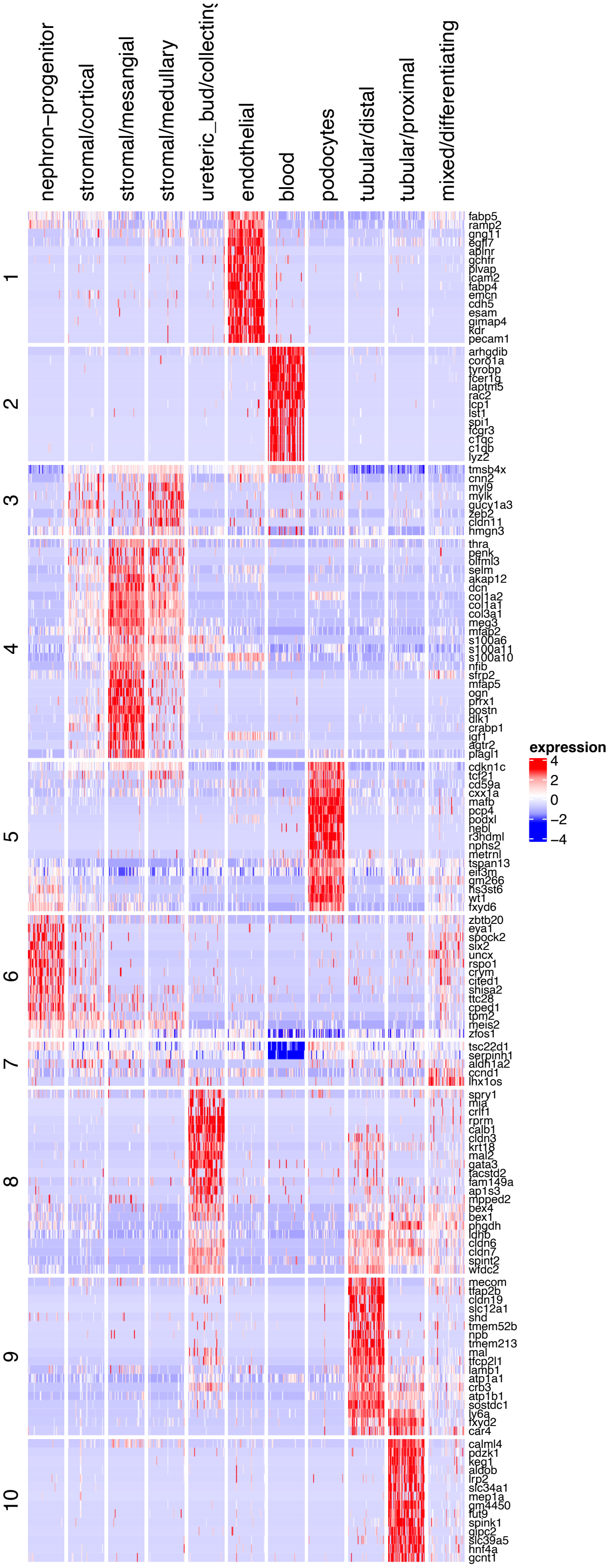

**Supplemental Figure 2**

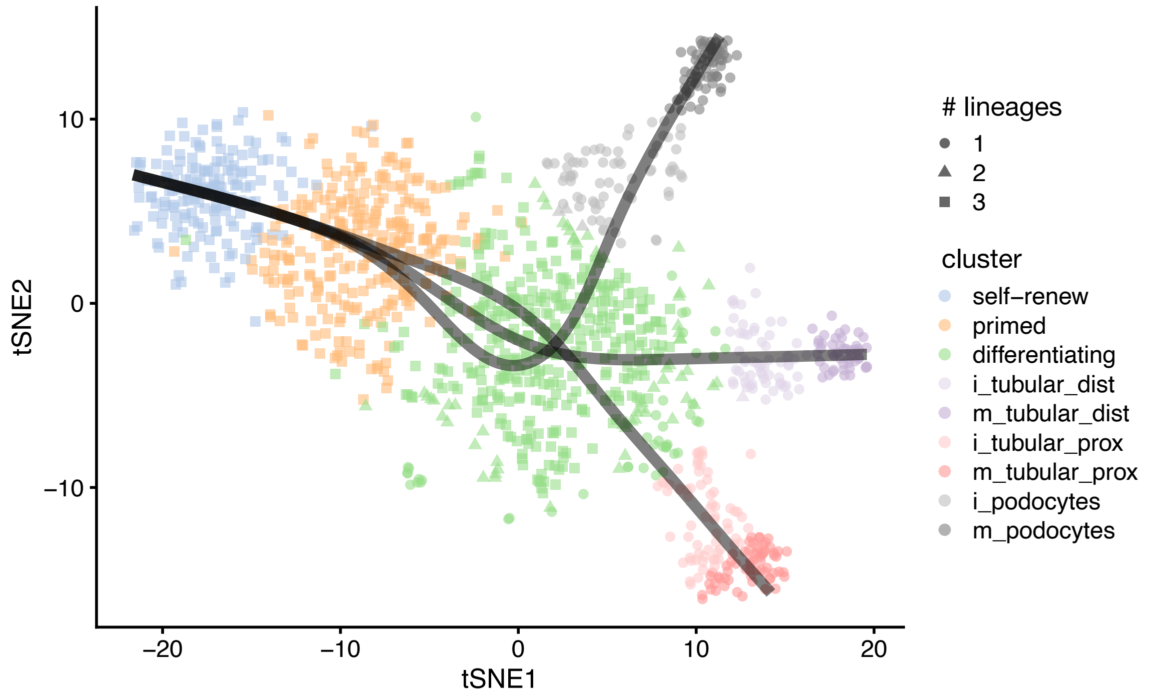

#### Supplemental Figure 3

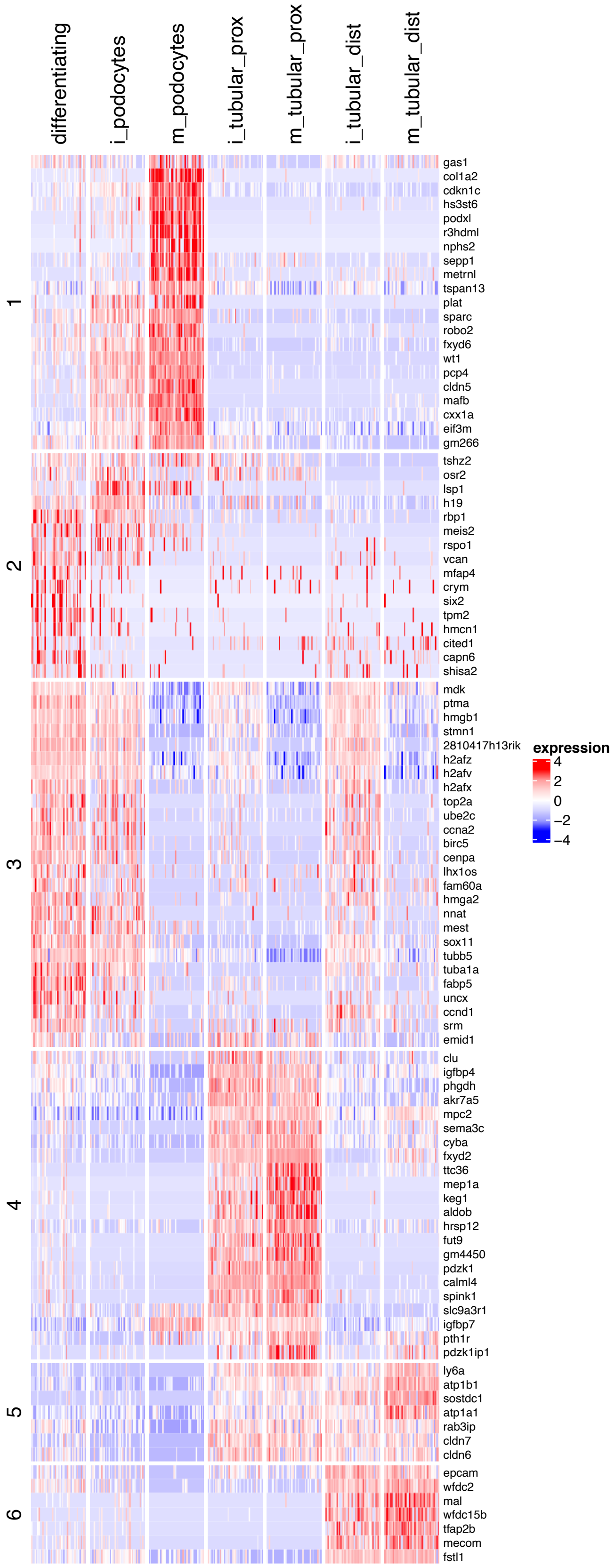

#### Supplemental Figure 4

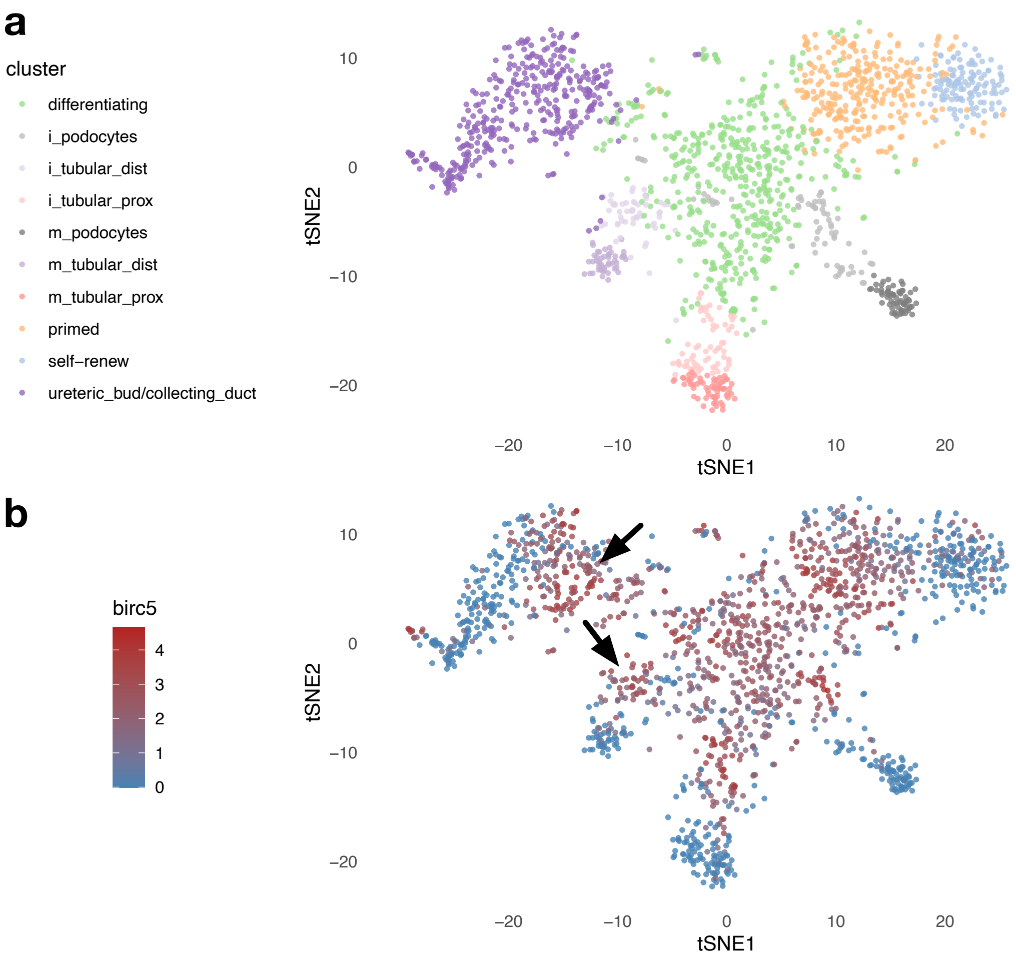

#### Supplemental Figure 5

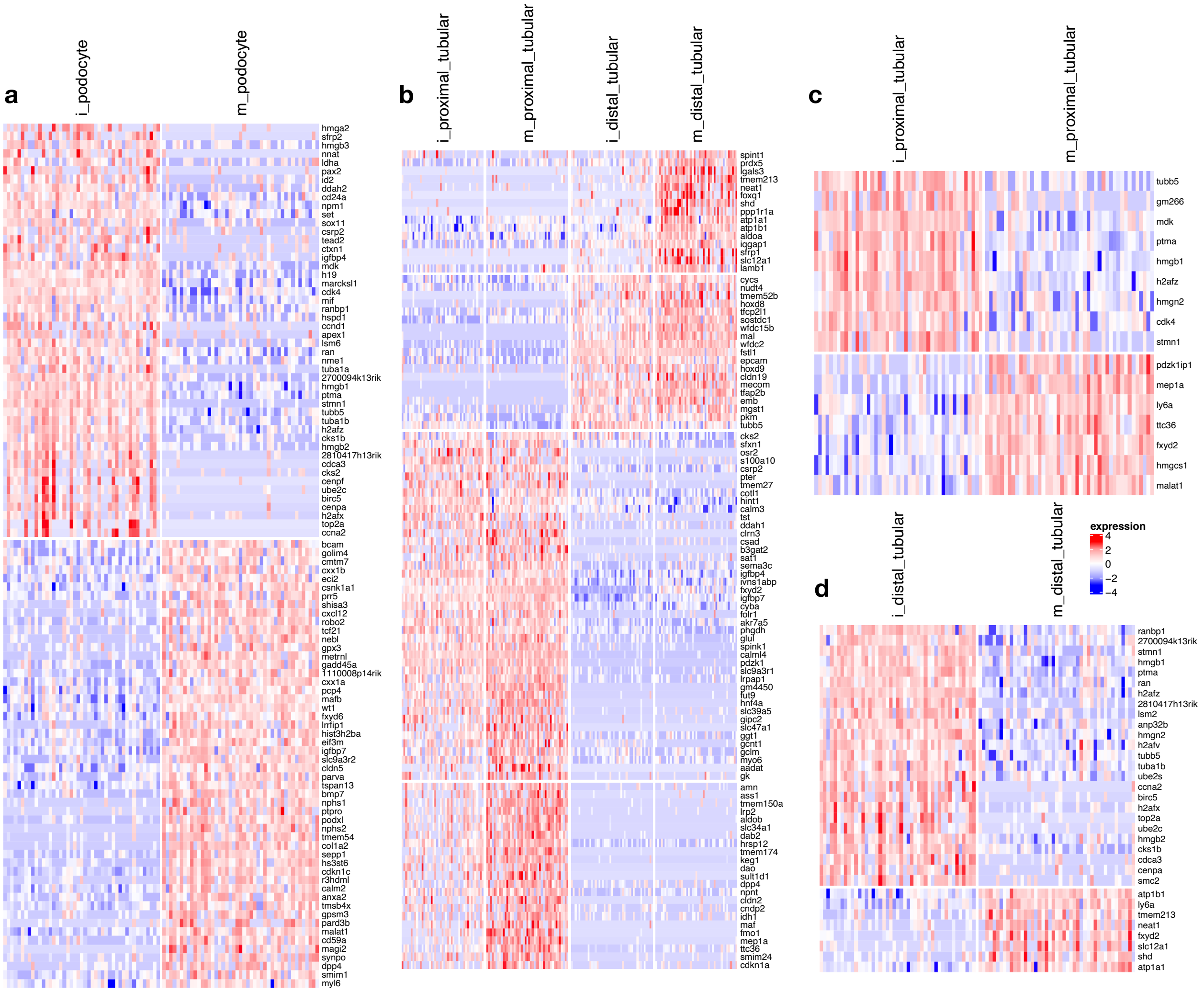
